# Single-molecule imaging reveals molecular coupling between transcription and DNA repair in live cells

**DOI:** 10.1101/515536

**Authors:** Han Ngoc Ho, Antoine van Oijen, Harshad Ghodke

**Author notes:** **Correspondence:** Harshad Ghodke, Molecular Horizons and School of Chemistry, University of Wollongong, Wollongong, New South Wales, 2522, Australia.

## Abstract

Actively transcribed genes are preferentially repaired in a conserved repair reaction known as transcription-coupled nucleotide excision repair^1–3^. During this reaction, stalled transcription elongation complexes at sites of lesions serve as a signal to trigger the assembly of nucleotide excision repair factors (reviewed in ref.^4,5^). In the model organism *Escherichia coli*, the transcription-repair coupling factor Mfd displaces the stalled RNA polymerase and hands-off the stall site to the nucleotide excision repair factors UvrAB for damage detection^6–9^. Despite *in vitro* evidence, it remains unclear how in live cells the stall site is faithfully handed over to UvrB from RNA polymerase and whether this handoff occurs via the Mfd-UvrA_2_-UvrB complex or via alternate reaction intermediates. Here, we visualise Mfd, the central player of transcription-coupled repair in actively growing cells and determine the catalytic requirements for faithful completion of the handoff during transcription-coupled repair. We find that the Mfd-UvrA_2_ complex is arrested on DNA in the absence of UvrB. Further, Mfd-UvrA_2_-UvrB complexes formed by UvrB mutants deficient in DNA loading and damage recognition, were also impaired in successful handoff. Our observations demonstrate that in live cells, the dissociation of Mfd is tightly coupled to successful loading of UvrB, providing a mechanism via which loading of UvrB occurs in a strand-specific manner during transcription-coupled repair.

---

DNA damage on the transcribed strand is repaired at a faster rate compared to lesions on the non-transcribed strand^1–3^. This enhanced transcription-coupled repair (TCR) is attributed to RNA polymerase (RNAP) acting as a damage sensor, followed by the Mfd-dependent recruitment of the nucleotide excision repair (NER) machinery^6^. A contemporary model for the TCR reaction is as follows: First, Mfd is recruited to the upstream edge of a stalled transcription elongation complex (TEC)^10^. This recruitment is accompanied by a release of Mfd’s auto-inhibition leading to the activation of its translocase activity that eventually disassembles RNAP, and a concomitant exposure of the UvrB homology module (BHM) to solution^7,11–14^. After disassembly of RNAP, Mfd continues to translocate on the DNA, and recruits the NER factors (UvrAB) in the vicinity of the lesion^9,15,16^. Despite successfully accommodating *in vitro* observations^9,15–17^, this model fails to answer several fundamental questions: How does the handoff complex direct the strand-specific loading of UvrB? Is this model valid inside living cells?

To address these gaps in understanding of the universally conserved TCR reaction, we employed single-molecule live-cell imaging to interrogate the handoff intermediate formed by fluorescently labelled Mfd (Mfd-YPet^18^) in its physiological context inside living *Escherichia coli*. Measurements of single molecules of Mfd-YPet in cells revealed that Mfd is recruited to DNA via pause/stalled TECs and that Mfd-YPet stays on DNA for 18 ± 1 s in wild-type cells^18^. In the absence of UvrA, the lifetime (τ_Mfd|Δ*uvrA*_) of the interaction of Mfd with stalled/paused RNAP on DNA increases to 29 ± 2 s ^18^. Having established that UvrA is necessary for completing the reaction with a lifetime of 18 s, we investigated whether UvrA alone was sufficient for promoting the dissociation of Mfd. To that end, we measured the lifetime of DNA-bound Mfd-YPet in cells lacking the downstream repair factor UvrB. First, we created a strain that expresses Mfd-YPet from the native chromosomal *mfd* locus while lacking the gene for UvrB (*mfd-YPet* Δ*uvrB*). Next, we immobilised these cells on a modified glass coverslip in a flow cell such that growing cells could be reliably imaged for several hours (Fig. 1a). The fluorescence of Mfd-YPet molecules manifested as a mixture of static foci arising from DNA-bound molecules, and cytosolic background arising from diffusive molecules (Fig. 1b). We first collected videos of cells expressing Mfd-YPet with continuous exposure times of 0.1 s (Fig. 1c, Extended Data Movie 1). Tracking foci of single Mfd-YPet molecules allows the determination of the rate of foci loss, which is a sum of photobleaching rate of the fluorescent protein and dissociation rate of Mfd-YPet (Fig. 1e, solid line).

**Figure 1.**
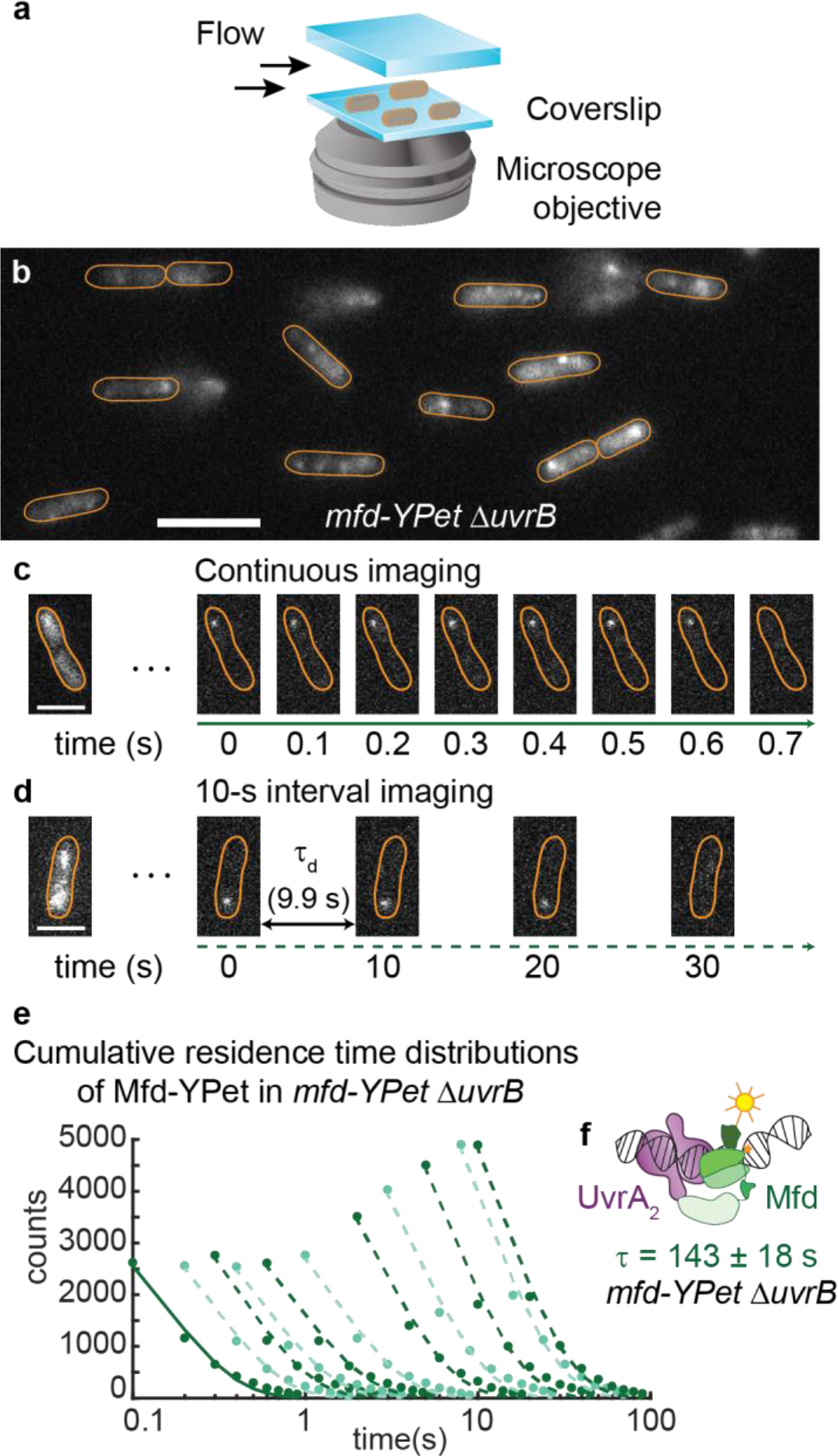
Measurements of Mfd-YPet kinetics of dissociation in cells lacking UvrB. **a**, Cells expressing fluorescent Mfd-YPet were grown to early exponential phase and loaded in a flow cell. Cells were imaged under constant supply of aerated growth medium for several hours. **b**, Representative fluorescence image of *mfd-YPet* Δ*uvrB* cells upon exposure to 514-nm light. Scale bar, 5 μm. Cell outlines (orange) are provided as a guide to the eye. **c**, Fluorescence data were collected in video format, with 0.1-s frames taken continuously. Due to fast photobleaching of the fluorescent protein YPet, continuous imaging hinders the observation of binding events on the second timescale. Scale bar, 2 μm. **d**, To extend the observation time window up to several minutes, images can be acquired with a dark interval (τ_d_) inserted between consecutive 0.1-s frames. Scale bar, 2 μm. **e**, Cumulative residence time distributions (CRTDs; circles) of Mfd-YPet in *mfd-YPet* Δ*uvrB* cells and the corresponding single-exponential fits (lines) obtained from continuous imaging (solid line) or interval imaging (dash lines) with a dark interval (τ_d_) between consecutive frames. τ_d_ increases from 0.1 s to 9.9 s (left to right, see Methods). **f**, Global fitting CRTDs in (**e**) yields a lifetime of 143 ± 18 s, suggesting Mfd (green) is arrested in complex with UvrA_2_ (purple) in cells lacking UvrB.

Here, the observation of long-lived binding events is limited due to the photobleaching of YPet upon continuous exposure to excitation photons. To extend the observation time window, we adopted interval imaging in which consecutive exposure times were spaced out by the addition of a dark time (τ_d_) (see ref.^19,20^ and Methods). We used a large set of distinct τ_d_ covering three orders of magnitude (0.1 s to 9.9 s) to ensure accurate measurements of binding lifetimes that last for seconds to several minutes inside cells (Fig. 1d-e). In cells lacking UvrB, the dissociation of Mfd-YPet is well described by single-exponential kinetics with a lifetime (τ_Mfd|Δ*uvrB*_) of 143 ± 18 s (Table 1, Extended Data Fig. 1). This lifetime is five times longer than in cells lacking UvrA (29 s), and eight times longer than in wild-type cells (18 s)^18^. Further, the lifetime of 143 s closely matches that of UvrA-YPet in *uvrA-YPet* Δ*uvrB* cells (97 ± 18 s) ^21^. Taken together, these results indicate that a highly stable DNA-bound Mfd-UvrA_2_ complex is formed in the absence of UvrB (Fig. 1f). Notably, UvrA can arrest the translocation of Mfd *in vitro*^9^.

**Table 1.** Lifetimes of Mfd-YPet in various genetic backgrounds. Mean ± s.d.

| **Derivatives of**<br>***mfd*-YPet** | **Slow lifetime**<br>**(s)** | **Percentage of the slow**<br>**population (%)** | **Fast**<br>**lifetime (s)** |
| --- | --- | --- | --- |
| $\Delta uvrB$ | 143 ± 18 | 100 | - |
| *uvrA*(K646A)<br>$\Delta uvrB$ | 26 ± 2 | 32 ± 2 | 1.1 ± 0.1 |
| *uvrA*(K646A) | 37 ± 3 | 100 | - |
| *uvrA*(K37A) $\Delta uvrB$ | 304 ± 69 | 100 | - |
| *uvrA*(K37A) | 52 ± 4 | 100 | - |
| *uvrB*($\Delta\beta$HG) | 188 ± 46 | 100 | - |
| *uvrB*(Y96A) | 70 ± 12 | 100 | - |
| $\Delta uvrB$ /pUvrB | 11.1 ± 0.7 | 26 ± 2 | 0.50 ± 0.08 |

Engagement of UvrA with UvrB and DNA is regulated by ATP hydrolysis at the two ATPase sites of UvrA^22–24^. To assess the role of ATP hydrolysis in the formation of the TCR handoff intermediate, we engineered *E. coli* strains that express either UvrA(K37A) (proximal ATPase mutant) or UvrA(K646A) (distal ATPase mutant) from the native *uvrA* locus in both *mfd-YPet* and *mfd-YPet* Δ*uvrB* backgrounds (Fig. 2a; Supplementary Method).

**Figure 2.**
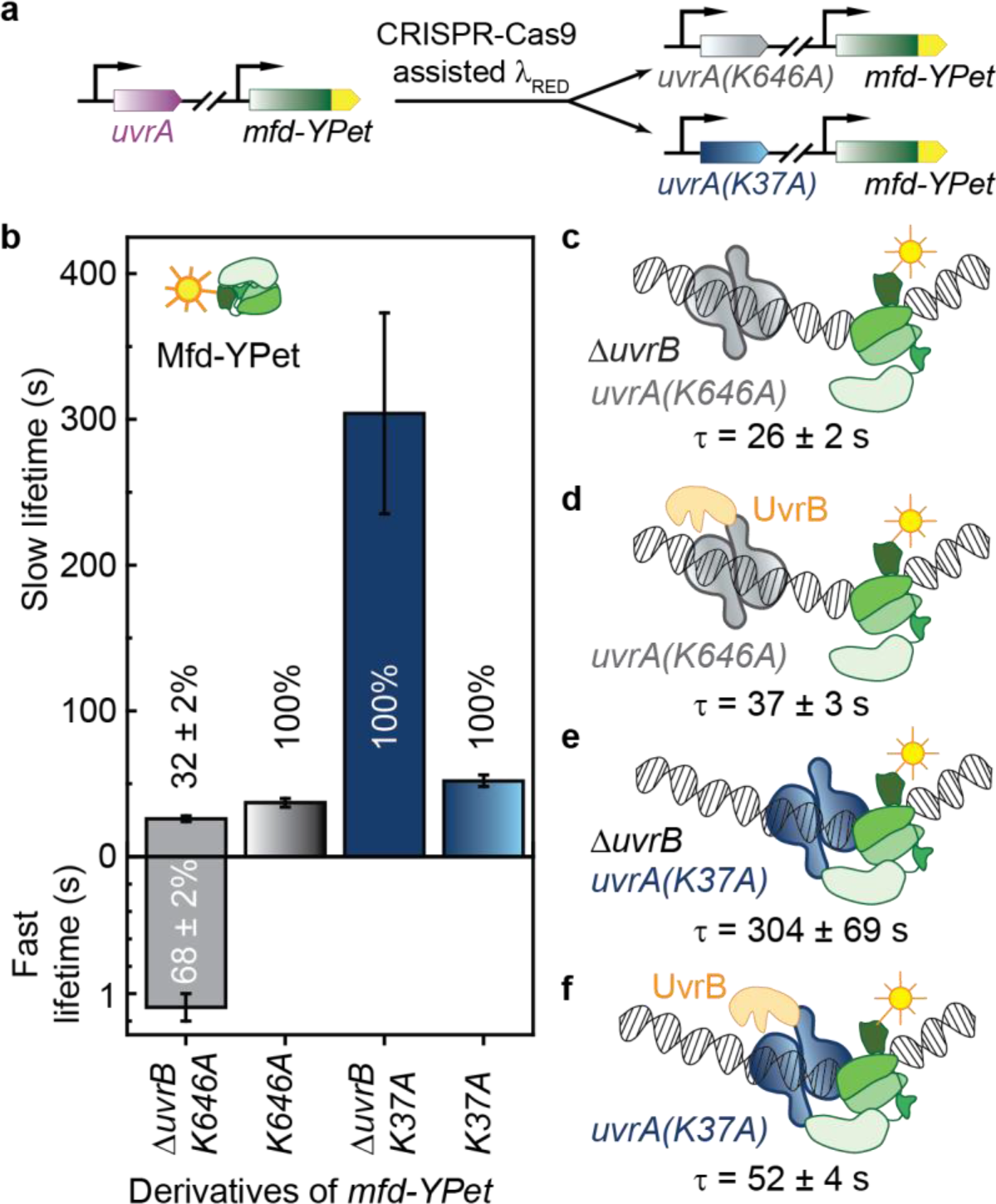
Measurements of Mfd-YPet kinetics of dissociation in cells expressing mutant UvrA deficient in ATP binding and hydrolysis. **a**, The wild-type *uvrA* allele was replaced by the mutant allele in *mfd-YPet* cells, so that mutant cells either express the distal ATPase mutant UvrA(K646A) or the proximal ATPase mutant UvrA(K37A) from the chromosome. **b**, Bar plots show lifetimes of DNA-bound Mfd-YPet in the corresponding genetic backgrounds. Where two kinetic sub-populations are detected, the fast lifetime is displayed in the lower panel. Percentages represent the amplitudes of kinetic sub-populations. Error bars are standard deviations from ten bootstrapped CRTDs (see Extended Data Fig. 2-3). **c**, Lifetime of Mfd-YPet in *mfd-YPet uvrA(K646A)* Δ*uvrB* (26 ± 2 s) is similar to that of Mfd-YPet in Δ*uvrA* cells (29 ± 2 s ^18^), suggesting the distal ATPase mutant (grey) is unable to interact with Mfd (green). **d**, Lifetime of Mfd-YPet in the presence of UvrA(K646A) and UvrB (orange) (37 ± 3 s) also resembles that of Mfd-YPet in Δ*uvrA* cells. **e**, Mfd-YPet is arrested in the presence of the proximal ATPase mutant UvrA(K37A) in cells lacking UvrB (lifetime of 304 ± 69 s). **f**, Mfd-YPet dissociates in 52 ± 4 s in the presence of UvrA(K37A) and UvrB, suggesting that ATP hydrolysis at the proximal site is required for promoting the dissociation of Mfd-YPet.

First, we measured the DNA-bound lifetimes of Mfd-YPet in cells expressing the UvrA(K646A) mutant. This mutant is severely defective in NER and TCR since it can load UvrB to 1% of the level of wild-type UvrA^11,22^. In the presence of UvrA(K646A), we detected two kinetic sub-populations of Mfd-YPet in cells lacking UvrB: a fast lifetime of 1.1 ± 0.1 (68 ± 2%) and a slow lifetime (τ_Mfd|*uvrA(K646A)* Δ*uvrB*_) of 26 ± 2 s (32 ± 2%) (Fig. 2b, Extended Data Fig. 2a-b). The fast dissociating sub-population (1.1 ± 0.1 s) may represent non-specific binding of Mfd-YPet to DNA^25^. Notably, the slow lifetime of 26 ± 2 s is comparable to that of Mfd-YPet in *mfd-YPet* Δ*uvrA* cells (29 ± 2 s)^18^. In the presence of UvrB, Mfd dissociated with a lifetime (τ_Mfd| *uvrA(K646A)*_) of 37 ± 3 s in cells carrying the UvrA(K646A) mutant (Fig. 2b; Extended Data Fig. 2c-d). Taken together, these results demonstrate that ATP hydrolysis at the distal ATPase site is critical for the formation of the Mfd-UvrA complex (Fig. 2c-d). A similar interpretation has been drawn in the case of the formation of the UvrAB complex^26^.

We then investigated the influence of the proximal ATPase on the lifetime of the Mfd-UvrA complex. In *mfd-YPet uvrA(K37A)* Δ*uvrB* cells, we found the lifetime of Mfd-YPet (τ_Mfd| *uvrA(K37A)* Δ*uvrB*_) to be 304 ± 69 s (Fig. 2b, Extended Data Fig. 3a-b), ten-fold longer than that of Mfd-YPet in Δ*uvrA* cells (τ_Mfd|Δ*uvrA*_ = 29 ± 2 s)^18^. The dissociation of Mfd-YPet in *mfd-YPet uvrA(K37A)* cells was six-fold faster in the presence of UvrB with a lifetime (τ_Mfd| *uvrA(K37A)*_) of 52 ± 4 s (Fig. 2b, Extended Data Fig. 3c-d). Since UvrA(K37A) can load UvrB with 10% efficiency compared to wild-type UvrA^22^, we propose that the complexes observed in our experiment likely represent a mixture of two populations – one that successfully loads UvrB like wild-type UvrA does and a population that is unable to load UvrB (Fig. 2e-f). Taken together, these measurements indicate that unlike UvrA(K646A) which fails to stably associate with Mfd, UvrA(K37A) can form the handoff complex, but fails to support the eviction of Mfd at rates comparable to that of wild-type UvrA. Consistent with this finding, UvrA(K37A) supports the preferential repair of the template strand *in vitro*^11^.

UvrA loads UvrB at the site of DNA damage during global genomic repair. In this step, single-stranded DNA is threaded in a cleft formed by the absolutely critical β-hairpin of UvrB^27,28^ (Extended Data Fig. 4a-b), followed by interrogation of the nucleobases mediated by UvrB’s cryptic helicase activity^29,30^. Since loading of UvrB promotes dissociation of UvrA from the damage surveillance complex^21^, we next investigated whether dissociation of Mfd from the handoff complex occurs upon loading of UvrB. We measured the residence time of Mfd-YPet in cells expressing β-hairpin mutants of UvrB from the native chromosomal locus^27,29,31^ (Extended Data Fig. 4c, Supplementary Method).

In cells carrying UvrB molecules lacking the β-hairpin (UvrB(ΔβHG)), the lifetime of Mfd (τ_Mfd|*uvrB(*Δβ*HG*_) was found to be 188 ± 46 s – an order of magnitude longer than in the wild-type UvrB background (Fig. 3a, Extended Data Fig. 4d-e). We then repeated these experiments in cells carrying the Y96A mutant of UvrB that can be loaded on DNA but fails to support damage verification. The lifetime of Mfd (τ_Mfd|*uvrB(Y96A)*_) was found to be 70 ± 12 s in this background (Fig. 3a, Extended Data Fig. 4f-g). Considering that both UvrB mutants retain the ability to form UvrAB-DNA complexes^29,31^, the simplest explanation is that the handoff complexes formed by Mfd-UvrA_2_ and mutant UvrB are impaired in evicting Mfd (Fig. 3b-c).

**Figure 3.**
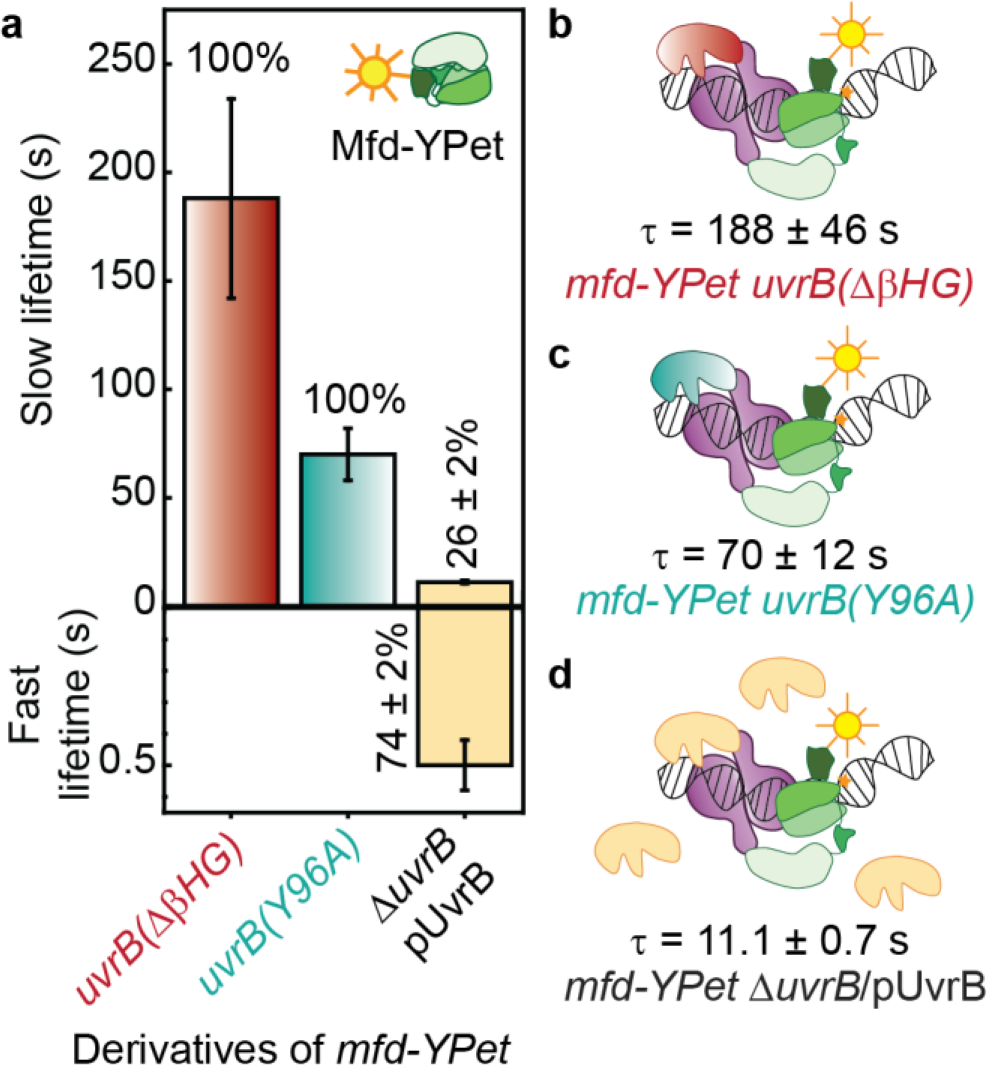
Measurements of Mfd-YPet kinetics of dissociation in cells expressing mutant UvrB deficient in DNA loading or over-expressing wild-type UvrB. **a**, Bar plots show lifetimes of DNA-bound Mfd-YPet in the corresponding genetic backgrounds. Where two kinetic sub-populations are detected, the fast lifetime is displayed in the lower panel. Percentages represent the amplitudes of kinetic sub-populations. Error bars are standard deviations from ten bootstrapped CRTDs (see Extended Data Fig. 4). **b**, Lifetime of Mfd-YPet in cells expressing mutant UvrB lacking the β-hairpin (red) was found to be 188 ± 46 s, demonstrating this mutant UvrB fails to promote the dissociation of Mfd-YPet (green). **c**, Similarly, mutant UvrB in which the absolutely conserve residue Y96 is replaced with alanine (cyan) also fails to promote the dissociation of Mfd-YPet (τ_Mfd| *uvrB(Y96A)*_ = 70 ± 12 s). **d**, Overexpression of UvrB (orange) leads to shorter lifetime of DNA-bound Mfd-YPet, compared to the constitutive level of UvrB (11.1 ± 0.7 s vs. 18 ± 1 s ^18^).

These results have three important implications: (1) since Mfd and UvrB do not physically interact, and UvrA is necessary for UvrB to be localised on DNA^32^, the stabilisation of Mfd on DNA in mutant UvrB backgrounds must occur via the Mfd-UvrA_2_-UvrB handoff complex. (2) UvrB does not simply compete off Mfd from UvrA_2_ complexes at the binding interface occupied by Mfd. (3) Binding of UvrB to the handoff complex is not sufficient for eviction of Mfd and UvrA from the handoff complex, successful engagement of DNA is a necessary requirement.

Since DNA repair enzymes must navigate the problem of target search, i.e., locating sparse DNA damage in a sea of undamaged DNA, we investigated whether the diffusion of UvrB to the site of the Mfd-UvrA_2_ complex limits the handoff rate. Elevated copy numbers of UvrB resulted in faster dissociation of Mfd (Fig. 3a,d; Extended Data Fig. 4h-k). In this case, the lifetime of Mfd (τ_Mfd| UvrB↑↑_) was 11.1 ± 0.7 s compared to the wild-type levels of UvrB (18 ± 1 s)^18^. These findings have broad implications – the rate limiting step of a reaction executed by a complex multicomponent machinery may be limited by concentrations of downstream factors. Since the expression of both UvrA and UvrB is repressed by LexA, SOS induction provides a means for elevating cellular concentrations of repair factors, consequently enhancing the rate of repair.

In this work, we show that the dissociation of Mfd is orchestrated by the ATPase activity of UvrA and is tightly coupled to the loading of UvrB in live bacterial cells. Considering that UvrB does not interact directly with Mfd^6^, the facilitated dissociation of Mfd observed in our experiments and *in vitro*^9^ indicates that UvrB exerts this influence via UvrA in a transient Mfd-UvrA_2_-UvrB handoff complex^6,9,16^. Importantly, experiments employing DNA loading mutants of UvrB demonstrate that the formation of this complex is not sufficient for promoting Mfd dissociation. In fact, the ability of UvrB to catalyse DNA loading is critical for the dissociation of Mfd from handoff complexes. Since dissociation of UvrA also requires loading of UvrB^6,21^, we propose that UvrA and Mfd dissociate from Mfd-UvrA_2_-UvrB handoff complexes in a single step. This is consistent with the original model proposed by Sancar lab, and with our observations that the residence times of Mfd and UvrA are similar. The simultaneous dissociation of Mfd and UvrA implies that UvrB is loaded in a strand-specific manner, providing an explanation for the observed preferential repair of the transcribed strand^16^. This model accommodates sequential recruitment of UvrA and UvrB to translocating Mfd as follows: Mfd is recruited to the site of a stalled TEC and is loaded in a strand-specific manner on the template strand carrying the putative DNA damage. This may be accompanied by translocation in the 3’-5’ direction which is subsequently arrested by binding of UvrA_2_^9^. ATPase-mediated structural changes of dimeric UvrA position UvrB in a conformation that favours loading of UvrB on the strand containing the DNA damage. Successful engagement of DNA by the β-hairpin of UvrB is then accompanied by dissociation of the Mfd-UvrA_2_ complex.

Faithful handoff of lesions to downstream factors is an inherent challenge faced by DNA repair machineries^33^. The observation that directional loading of the damage verification machinery serves as a molecular switch to trigger the dissociation of upstream damage recognition factors provides an elegant solution to this challenge. Since the eukaryotic CSB is a functional (and structural) homolog of Mfd^34^, it will be interesting to see whether coupled, directional loading of the repair machinery is a fundamentally conserved feature of the transcription-coupled repair reaction.

## METHODS

### Construction of strains and plasmid

All strains used in this study were derivatives of *Escherichia coli* K-12 MG1655 *mfd-YPet*^18^ (see Extended Data Table 1). Point mutations in chromosomal *uvrA* or *uvrB* allele were introduced by CRISPR-Cas9 assisted λ Red recombination^35,36^ (see Supplementary Method and Extended Data Table 2). Knock-outs of *uvrB* allele were aided by P1 transduction.

pUvrB was created by sub-cloning the *uvrB* promoter and *uvrB* gene (synthetic dsDNA, IDT, Illinois, US) into pJM1071 (a gift from Woodgate lab)^37^ at *NdeI* and *Xho*I sites. The promoter sequence was identified as 130 nucleotides directly upstream of the *uvrB* gene in the *E. coli* chromosome^38^. Mutant alleles on the chromosome and pUvrB were sequenced on both strands prior to use.

### Cell culture for imaging

Cells were imaged in quartz-top flow cells as described previously^18^. Cells were grown in 500 μL of EZ-rich defined media (Teknova, CA, US), supplemented with 0.2% (v/v) glucose in 2 mL microcentrifuge tubes at 30 °C. For experiments involving plasmid-expressed UvrB, spectinomycin (50 μg per mL) was added to the growth media. Cells in early exponential phase were loaded in flow cells at 30 °C, followed by a constant supply of aerated EZ-rich defined media at a rate of 30 μL per min, using a syringe pump (Adelab Scientific, Australia).

### Live-cell imaging

Single-molecule fluorescence imaging was carried out with a custom-built microscope as previously described^18^. Briefly, the microscope comprised a Nikon Eclipse Ti body, a 1.49 NA 100× objective, a 514-nm Sapphire LP laser (Coherent) operating at a power density of 71 W.cm^−2^, an ET535/30m emission filter (Chroma) and a 512 × 512 pixel^2^ EM-CCD camera (either Photometrics Evolve or Andor iXon 897). The microscope operated in near-TIRF illumination^39^ and was controlled using NIS-Elements (Nikon).

Fluorescence images were acquired in time-series format with 0.1-s frames. Each video acquisition contained two phases. The first phase aimed to lower background signal by continuous illuminating, causing most of the fluorophores to photo-bleach or to assume a dark state. The second phase (single-molecule phase) is when single molecules can be reliably tracked on a low background signal. In the second phase, consecutive frames were acquired continuously or with a dark time (τ_d_).

### Image analysis

Image analysis was performed in Fiji^40^, using the Single Molecule Biophysics plugins (available at https://github.com/SingleMolecule/smb-plugins), and MATLAB. First, raw data were converted to TIF format, following by background correction and image flattening as previously described^18^. Next, foci were detected in the reactivation phase by applying a discoidal average filter (inner radius of one pixel, outer radius of three pixels), then selecting pixels above the intensity threshold. Foci detected within 3-pixel radius (318 nm) in consecutive frames were considered to belong to the same binding event.

### Interval imaging for dissociation kinetics measurements

Interval imaging was performed as described previously^18^. The photobleaching phase contained 50 continuous 0.1-s frames. In phase II, 100 0.1-s frames were collected either continuously or with a delay time (τ_d_) ranging from τ_d_ = (0.1, 0.2, 0.3, 0.5, 0.9, 1.9, 2.9, 4.9, 7.9, 9.9). In each experiment, videos with varying τd were acquired. Foci were detected using a relative intensity threshold of 8 above the background. Depending on the construct being imaged, between 5-11 repeats of each experiment were collected for each strain. Cumulative residence time distribution (CRTDs) of binding events detected in all data sets were generated for each interval. Lifetimes of DNA-bound Mfd-YPet were determined by globally fitting bootstrapped CRTDs across all intervals using least-squares trust-region reflective algorithms as described in ref.^20^. The iteration terminated when a tolerance of 10^−6^ was reached. Dissociation rates and the corresponding lifetimes were means of ten bootstrapped samples, each derived from randomly selecting 80% of the compiled binding events (Extended Data Table 3).

## ACKNOWLEDGEMENTS

We thank Lidia G. Alvarez for collecting microscope data during the early stage of the project. H.G. acknowledges the University of Wollongong and the Faculty of Science, Medicine and Health for funding of the Andor iXon 897 used in this work. A.M. van Oijen would like to acknowledge support by the Australian Research Council (DP150100956 and FL140100027).

## AUTHOR CONTRIBUTIONS

Construct creation: H.N.H.; Data curation: H.N.H.; Data analysis: H.N.H. and H.G.; Software: H.N.H. and H.G.; Writing—Original draft: H.N.H. and H.G.; Writing—review and editing: H.N.H., H.G. and A.M.v.O.; Conceptualization: H.G.; Supervision: H.G. and A.M.v.O.

## MATERIALS AND CORRESPONDENCE

Harshad Ghodke, Molecular Horizons and School of Chemistry and Molecular Bioscience, University of Wollongong, Wollongong, New South Wales, 2522, Australia

## REFERENCES

1. Mellon, I., Spivak G. & Hanawalt, P. C. Selective removal of transcription-blocking DNA damage from the transcribed strand of the mammalian DHFR genes. Cell 51, 241–249 (1987).

2. Mellon I. & Hanawalt P. C. Induction of the Escherichia coli lactose operon selectively increases repair of its transcribed DNA strand. Nature 342, 95–98, doi:10.1038/342095a0 (1989).

3. Smerdon M. J. & Thoma F. Site-specific DNA repair at the nucleosome level in a yeast minichromosome. Cell 61, 675–684 (1990).

4. Hanawalt P. C. & Spivak G. Transcription-coupled DNA repair: two decades of progress and surprises. Nat Rev Mol Cell Biol 9, 958–970, doi:10.1038/nrm2549 (2008).

5. Gregersen L. H. & Svejstrup J. Q. The Cellular Response to Transcription-Blocking DNA Damage. Trends Biochem Sci 43, 327–341, doi:10.1016/j.tibs.2018.02.010 (2018).

6. Selby C. P. & Sancar A. Molecular mechanism of transcription-repair coupling. Science 260, 53–58 (1993).

7. Deaconescu A. M. et al. Structural Basis for Bacterial Transcription-Coupled DNA Repair. Cell 124, 507–520 (2006).

8. Howan K. et al. Initiation of transcription-coupled repair characterized at single-molecule resolution. Nature 490, 431–434 (2012).

9. Fan J., Leroux-Coyau, M., Savery N. J. & Strick T. R. Reconstruction of bacterial transcription-coupled repair at single-molecule resolution. Nature 536, 234–237, doi:10.1038/nature19080 (2016).

10. Park, J.-S., Marr M. T. & Roberts J. W. E. coli Transcription Repair Coupling Factor (Mfd Protein) Rescues Arrested Complexes by Promoting Forward Translocation. Cell 109, 757–767 (2002).

11. Manelyte L., Kim Y. I., Smith A. J., Smith R. M. & Savery N. J. Regulation and rate enhancement during transcription-coupled DNA repair. Mol Cell 40, 714–724, doi:10.1016/j.molcel.2010.11.012 (2010).

12. Smith A. J., Szczelkun M. D. & Savery N. J. Controlling the motor activity of a transcription-repair coupling factor: autoinhibition and the role of RNA polymerase. Nucleic Acids Res 35, 1802–1811, doi:10.1093/nar/gkm019 (2007).

13. Murphy M. N. et al. An N-terminal clamp restrains the motor domains of the bacterial transcription-repair coupling factor Mfd. Nucleic Acids Res 37, 6042–6053, doi:10.1093/nar/gkp680 (2009).

14. Deaconescu A. M., Sevostyanova A., Artsimovitch I. & Grigorieff N. Nucleotide excision repair (NER) machinery recruitment by the transcription-repair coupling factor involves unmasking of a conserved intramolecular interface. Proc Natl Acad Sci U S A 109, 3353–3358, doi:10.1073/pnas.1115105109 (2012).

15. Graves E. T. et al. A dynamic DNA-repair complex observed by correlative single-molecule nanomanipulation and fluorescence. Nat Struct Mol Biol 22, 452–457 (2015).

16. Haines N. M., Kim, Y.-I. T., Smith A. J. & Savery N. J. Stalled transcription complexes promote DNA repair at a distance. Proceedings of the National Academy of Sciences 111, 4037–4042, doi:10.1073/pnas.1322350111 (2014).

17. Howan K. et al. Initiation of transcription-coupled repair characterized at single-molecule resolution. Nature 490, 431–434, doi:10.1038/nature11430 (2012).

18. Ho H. N., van Oijen, A. M. & Ghodke H. The transcription-repair coupling factor Mfd associates with RNA polymerase in the absence of exogenous damage. Nat Commun 9, 1570, doi:10.1038/s41467-018-03790-z (2018).

19. Gebhardt J. C. et al. Single-molecule imaging of transcription factor binding to DNA in live mammalian cells. Nat Methods 10, 421–426, doi:10.1038/nmeth.2411 (2013).

20. Ho H. N., Zalami D., Köhler, J., van Oijen, A.M., Ghodke H. Identifying multiple kinetic populations of DNA binding proteins in live cells using single-molecule fluorescence imaging. bioRxiv, doi:https://doi.org/10.1101/509620 (2019).

21. Ghodke H., Ho, H.N., van Oijen, A.M. Single-molecule live-cell imaging reveals parallel pathways of prokaryotic nucleotide excision repair. bioRxiv (2019).

22. Myles G. M., Hearst J. E. & Sancar A. Site-specific mutagenesis of conserved residues within Walker A and B sequences of Escherichia coli UvrA protein. Biochemistry 30, 3824–3834 (1991).

23. Thiagalingam S. & Grossman L. Both ATPase sites of Escherichia coli UvrA have functional roles in nucleotide excision repair. J Biol Chem 266, 11395–11403 (1991).

24. Wagner K., Moolenaar G. F. & Goosen N. Role of the two ATPase domains of Escherichia coli UvrA in binding non-bulky DNA lesions and interaction with UvrB. DNA Repair (Amst) 9, 1176–1186, doi:10.1016/j.dnarep.2010.08.008 (2010).

25. Selby C. P. & Sancar A. Structure and function of transcription-repair coupling factor. I. Structural domains and binding properties. J Biol Chem 270, 4882–4889 (1995).

26. Stracy M. et al. Single-molecule imaging of UvrA and UvrB recruitment to DNA lesions in living Escherichia coli. Nat Commun 7, 12568, doi:10.1038/ncomms12568 (2016).

27. Truglio J. J. et al. Structural basis for DNA recognition and processing by UvrB. Nat Struct Mol Biol 13, 360–364, doi:10.1038/nsmb1072 (2006).

28. Moolenaar G. F., Hoglund L. & Goosen N. Clue to damage recognition by UvrB: residues in the beta-hairpin structure prevent binding to non-damaged DNA. EMBO J 20, 6140–6149, doi:10.1093/emboj/20.21.6140 (2001).

29. Skorvaga M. et al. Identification of residues within UvrB that are important for efficient DNA binding and damage processing. J Biol Chem 279, 51574–51580, doi:10.1074/jbc.M409266200 (2004).

30. Caron P. R. & Grossman L. Involvement of a cryptic ATPase activity of UvrB and its proteolysis product, UvrB* in DNA repair. Nucleic Acids Res 16, 10891–10902 (1988).

31. Skorvaga M., Theis K., Mandavilli B. S., Kisker C. & Van Houten, B. The beta-hairpin motif of UvrB is essential for DNA binding, damage processing, and UvrC-mediated incisions. J Biol Chem 277, 1553–1559, doi:10.1074/jbc.M108847200 (2002).

32. Kad N. M., Wang H., Kennedy G. G., Warshaw D. M. & Van Houten, B. Collaborative dynamic DNA scanning by nucleotide excision repair proteins investigated by single-molecule imaging of quantum-dot-labeled proteins. Mol Cell 37, 702–713, doi:10.1016/j.molcel.2010.02.003 (2010).

33. Liu J., Lee J. B. & Fishel R. Stochastic Processes and Component Plasticity Governing DNA Mismatch Repair. J Mol Biol 430, 4456–4468, doi:10.1016/j.jmb.2018.05.039 (2018).

34. Xu J. et al. Structural basis for the initiation of eukaryotic transcription-coupled DNA repair. Nature 551, 653–657, doi:10.1038/nature24658 (2017).

35. Jiang W., Bikard D., Cox D., Zhang F. & Marraffini L. A. RNA-guided editing of bacterial genomes using CRISPR-Cas systems. Nat Biotechnol 31, 233–239, doi:10.1038/nbt.2508 (2013).

36. Pyne M. E., Moo-Young, M., Chung D. A. & Chou C. P. Coupling the CRISPR/Cas9 System with Lambda Red Recombineering Enables Simplified Chromosomal Gene Replacement in Escherichia coli. Appl Environ Microbiol 81, 5103–5114, doi:10.1128/AEM.01248-15 (2015).

37. Churchward G., Belin D. & Nagamine Y. A pSC101-derived plasmid which shows no sequence homology to other commonly used cloning vectors. Gene 31, 165–171 (1984).

38. Sancar G. B., Sancar A., Little J. W. & Rupp W. D. The uvrB gene of Escherichia coli has both lexA-repressed and lexA-independent promoters. Cell 28, 523–530 (1982).

39. Tokunaga M., Imamoto N. & Sakata-Sogawa, K. Highly inclined thin illumination enables clear single-molecule imaging in cells. Nat Methods 5, 159–161, doi:10.1038/nmeth1171 (2008).

40. Schindelin J. et al. Fiji: an open-source platform for biological-image analysis. Nat Methods 9, 676–682, doi:10.1038/nmeth.2019 (2012).

