## Supplementary Information for "Single-molecule imaging reveals molecular coupling between transcription and DNA repair in live cells"

#### Supplementary Figures

**Extended Data Figure 1:** The $k_{eff}\tau_{tl}$ plot for *mfd-YPet ΔuvrB* (solid line) indicates a single slowly dissociating species. For comparison, two simulated $k_{eff}\tau_{tl}$ plots describing a single dissociating species with $k_{off}$ of 3.5x10$^{-2}$ s$^{-1}$ ($\tau_1$ = 29 s; blue dashed curve) or 7x10$^{-3}$ s$^{-1}$ ($\tau_2$ = 143 s; grey dashed curve) are provided. Shaded error bands represent standard deviations from ten bootstrapped samples. Simulations were performed with custom-written MATLAB codes as described in ref.[1].

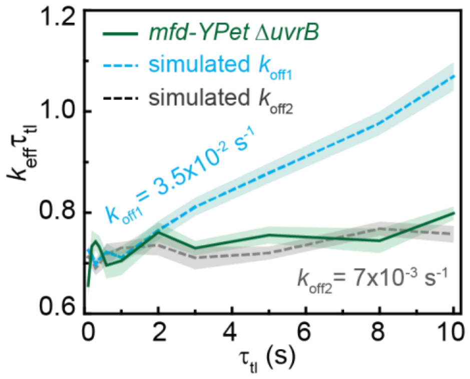

**Extended Data Figure 2:** Measurements of Mfd-YPet kinetics of dissociation in cells expressing the distal ATPase mutant UvrA(K646A).

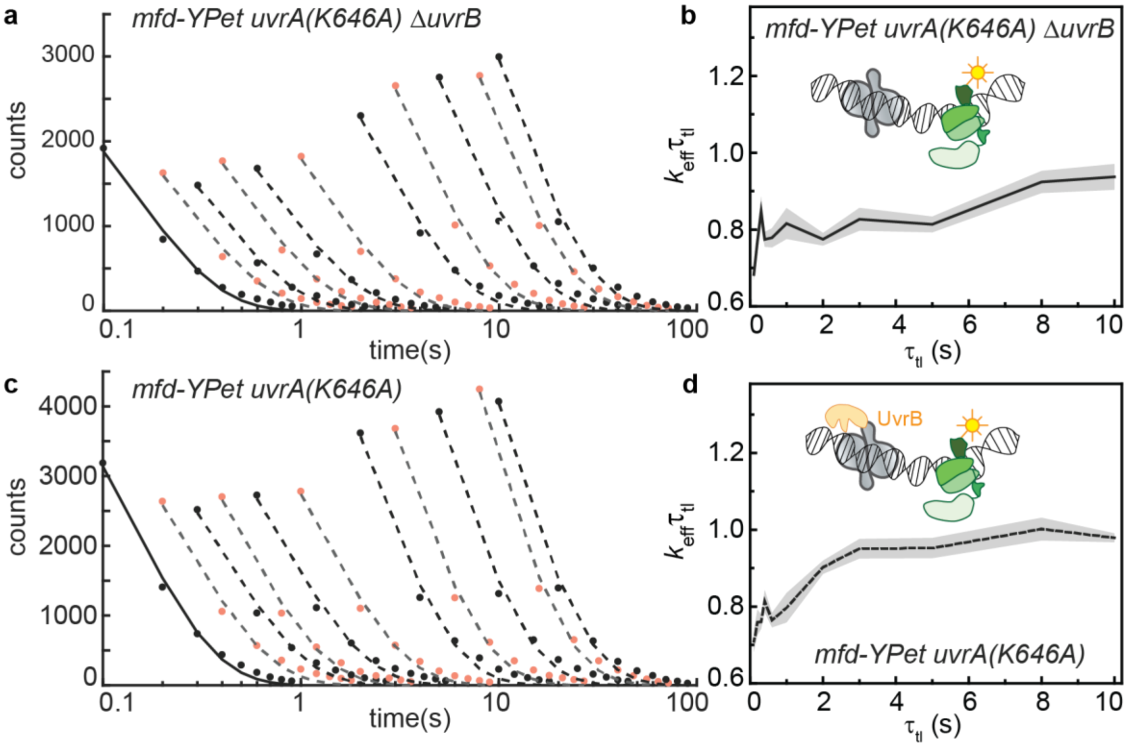


**a**. Cumulative residence time distributions (CRTDs, circles) obtained from interval imaging of Mfd-YPet in *mfd-YPet uvrA(K646A) ΔuvrB* cells. Lines are mono-exponential fits to CRTDs.

**b**. The $k_{eff}\tau_{tl}$ plot obtained from fitting CRTDs of Mfd-YPet in *mfd-YPet uvrA(K646A) ΔuvrB* cells. Shaded error bands are standard deviations from ten bootstrapped samples. Cartoon (inset) illustrates the inability of UvrA(K646A) (grey) to interact with Mfd-YPet (green).

**c**. CRTDs (circles) obtained from interval imaging of Mfd-YPet in *mfd-YPet uvrA(K646A)* cells. Lines are mono-exponential fits to CRTDs.

**d**. The $k_{eff}\tau_{tl}$ plot obtained from fitting CRTDs of Mfd-YPet in *mfd-YPet uvrA(K646A)* cells. Shaded error bands are standard deviations from ten bootstrapped samples. Cartoon (inset) illustrates the inability of UvrA(K646A)-UvrB (grey and orange) complex to interact with Mfd-YPet (green).

**Extended Data Figure 3:** Measurements of Mfd-YPet kinetics of dissociation in cells expressing the proximal ATPase mutant UvrA(K37A).

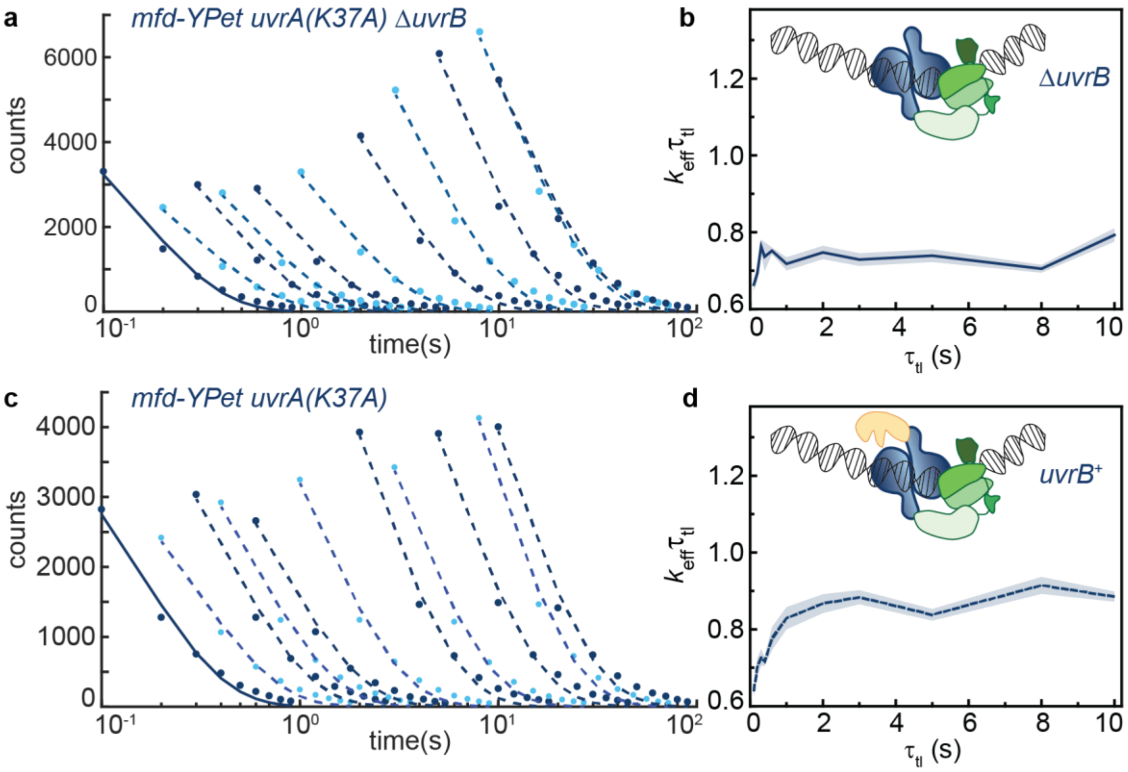


**a**. Cumulative residence time distributions (CRTDs, circles) obtained from interval imaging of Mfd-YPet in *mfd-YPet uvrA(K37A) ΔuvrB* cells. Lines are mono-exponential fits to CRTDs.

**b**. The $k_{eff}\tau_{tl}$ plot obtained from fitting CRTDs of Mfd-YPet in *mfd-YPet uvrA(K37A) ΔuvrB* cells. Shaded error bands are standard deviations from ten bootstrapped samples. Cartoon (inset) illustrates the arrested complex formed by UvrA(K37A) (blue) and Mfd (green).

**c**. CRTDs (circles) obtained from interval imaging of Mfd-YPet in *mfd-YPet uvrA(K37A)* cells. Lines are mono-exponential fits to CRTDs.

**d**. The $k_{eff}\tau_{tl}$ plot obtained from fitting CRTDs of Mfd-YPet in *mfd-YPet uvrA(K37A)* cells. Shaded error bands are standard deviations from ten bootstrapped samples. Cartoon (inset) illustrates the impaired handoff complex formed by UvrB (orange), UvrA(K37A) (blue) and Mfd (green).

**Extended Data Figure 4:** Measurements of Mfd-YPet kinetics of dissociation in cells expressing mutant UvrB deficient in DNA loading or over-expressing wild-type UvrB.

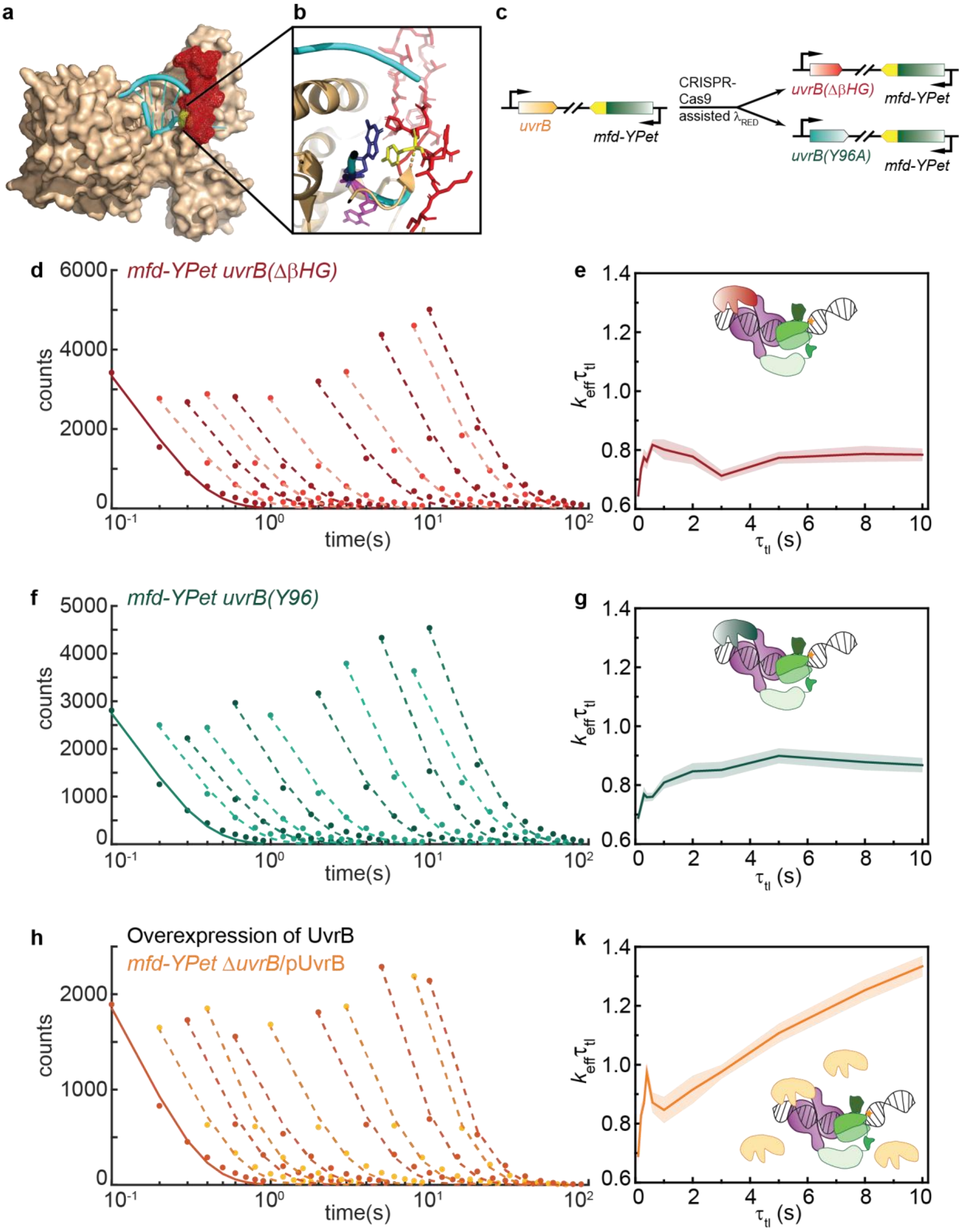

**a**. Crystal structure of *Bacillus caldotenax* UvrB (brown) bound to a DNA hairpin (cyan) (PDB ID: 2FDC)[2]. The β-hairpin of UvrB is highlighted in red.

**b**. Zoomed-in view of the tyrosine residue Y96 (yellow) at the base of the β-hairpin (red). The tyrosine residue stabilises the DNA by forming π-stacking interactions with a guanine (blue) while a neighbouring base (magenta) is flipped out into a hydrophobic pocket of UvrB.

**c**. UvrB mutants deficient in loading are expressed from the chromosome of *mfd-YPet* cells, where the native *uvrB* gene has been edited using CRISPR-Cas9 assisted λ Red recombination.

**d**. CRTDs (circles) obtained from interval imaging of Mfd-YPet in *mfd-YPet uvrB(*ΔβHG*)* cells. Lines are mono-exponential fits to CRTDs.

**e**. The $k_{\mathrm{eff}}\tau_{\mathrm{tl}}$ plot obtained from fitting CRTDs of Mfd-YPet in *mfd-YPet uvrB(*ΔβHG*)* cells. Shaded error bands are standard deviations from ten bootstrapped samples. Cartoon (inset) illustrates the impaired handoff complex formed by UvrB(ΔβHG) (red), UvrA (purple) and Mfd (green).

**f**. CRTDs (circles) obtained from interval imaging of Mfd-YPet in *mfd-YPet uvrB(Y96A)* cells. Lines are mono-exponential fits to CRTDs.

**g**. The $k_{\mathrm{eff}}\tau_{\mathrm{tl}}$ plot obtained from fitting CRTDs of Mfd-YPet in *mfd-YPet uvrB(Y96A)* cells. Shaded error bands are standard deviations from ten bootstrapped samples. Cartoon (inset) illustrates the impaired handoff complex formed by UvrB(Y96A) (cyan), UvrA (purple) and Mfd (green).

**h**. CRTDs (circles) obtained from interval imaging of Mfd-YPet in *mfd-YPet* Δ*uvrB*/pUvrB cells, in which UvrB is over-expressed from the pUvrB plasmid. Lines are mono-exponential fits to CRTDs.

**k**. The $k_{\mathrm{eff}}\tau_{\mathrm{tl}}$ plot obtained from fitting CRTDs of Mfd-YPet in *mfd-YPet* Δ*uvrB*/pUvrB cells. Shaded error bands are standard deviations from ten bootstrapped samples. Cartoon (inset) illustrates the facilitated dissociation of Mfd in the context of the UvrB-$UvrA_2$-Mfd handoff complex.

### Supplementary Tables

**Extended Data Table 1:** Bacterial strains. All strains are in *E. coli* K-12 MG1655 background.

| Strain/genotypes | Source/Technique |
|---|---|
| MG1655 *mfd-YPet* | This laboratory |
| *uvrB::kanR* | This study/ λ Red recombination |
| *mfd-YPet* Δ*uvrB* | This study/ P1 transduction |
| *mfd-YPet* Δ*uvrB*/pUvrB | This study |
| *mfd-YPet uvrA(K646A)* | This study/ CRISPR-Cas9 assisted λ Red recombination |
| *mfd-YPet uvrA(K646A) uvrB::kanR* | This study/ P1 transduction |
| *mfd-YPet uvrA(K37A)* | This study/ CRISPR-Cas9 assisted λ Red recombination |
| *mfd-YPet uvrA(K37A) uvrB::kanR* | This study/ P1 transduction |
| *mfd-YPet uvrB(ΔβHG)* | This study/ CRISPR-Cas9 assisted λ Red recombination |
| *mfd-YPet uvrB(Y96A)* | This study/ CRISPR-Cas9 assisted λ Red recombination |

**Extended Data Table 2:** Oligonucleotides used for colony PCR, λ Red recombination and cloning.

| Oligo names | Sequence |
|---|---|
| *Colony PCR for Cas9 verification* | |
| dCas9dL5_303_F | CAGACCGCCACAGTATCAAA |
| pCas9_6700_R | GGAAGGTATCCGACTGCTG |
| *Cloning of pCRISPR variants* | |
| pCRISPR_UvrA_K646A_S | AAA CGT TAA TCA GCG TCG ATT TAC G |
| pCRISPR_UvrA_K646A_AS | AAA ACG TAA ATC GAC GCT GAT TAA C |
| pCRISPR_UvrA_K37A_S | AAA CGT GAC CGG GCT TTC GGG TTC G |
| pCRISPR_UvrA_K37A_AS | AAA ACG AAC CCG AAA GCC CGG TCA C |
| pCRISPR_UvrB_Y96A_S | AAA CCC TAC TAC GAC TAC TAT CAG CG |
| pCRISPR_UvrB_Y96A_AS | AAA ACG CTG ATA GTA GTC GTA GTA GG |
| *Recombineric ssDNA* | |
| UvrA_K646A_ ssDNA | GGC GTT GGG CAA TCG GGA ACA GTG TGT CGT TAA TCA GCG TCG ACG CAC CGG AAC CTG AAA CCC CGG TGA TGC AGG TAA ACA GAC CCA |
| UvrA_K37A_ ssDNA | GCT GCC CTT CGG CAT ATA AGG TGT CGA AAG CGA GCG AGG ACG CGC CGC TAC CCG AAA GCC CGG TCA CGA CAA TGA GCT TGT CGC GGG |
| UvrB_Y96A_ ssDNA | CAA TGA AAG TGT CGG AAC TCG GTA CAT AGG CTT CCG GCT GTG CGT AGT CGT AGT AGG AAA CGA AAT ATT CCA CCG CGT TTT CC |
| UvrB_ΔβHG_ ssDNA | CAA TAT GTT CGT TAA CCG AGG CAT CTT TCT CAA TGA AAG TGC CAT AAT AGT CGT AGT AGG AAA CGA AAT ATT CCA CCG CGT TTT CCG |

**Extended Data Table 3:** Global fitting outputs for measurements of lifetime of Mfd-YPet in various genetic backgrounds. Bootstrapped CRTDs were fitted two single- and bi-exponential models (model 1 and 2 respectively). Selection of model was carried out as in ref.[1]. The chosen model and fitting outcomes are highlighted. Errors are standard deviations from ten bootstrapped CRTDs.

| Derivatives of *mfd-YPet* | Model | $k_b$ ± Error ($s^{-1}$) | $\tau_1$ ± Error (s) | *B* ± Error (%) | $\tau_2$ ± Error (s) |
|---|---|---|---|---|---|
| *ΔuvrB* | **1** | **7.1 ± 0.1** | **143 ± 18** | **100** | **-** |
| | 2 | 6.2 ± 0.1 | 1000 | 41 ± 2 | 5.5 ± 0.4 |
| *uvrA(K646A) ΔuvrB* | 1 | 7.5 ± 0.1 | 54 ± 8 | 100 | - |
| | **2** | **5.9 ± 0.1** | **26 ± 2** | **32 ± 2** | **1.1 ± 0.1** |
| *uvrA(K646A)* | **1** | **7.9 ± 0.1** | **37 ± 3** | **100** | **-** |
| | 2 | 6.9 ± 0.1 | 1000 | 26 ± 2 | 5.4 ± 0.2 |
| *uvrA(K37A) ΔuvrB* | **1** | **7.2 ± 0.1** | **304 ± 69** | **100** | **-** |
| | 2 | 6.0 ± 0.1 | 1000 | 40 ± 2 | 4.7 ± 0.2 |
| *uvrA(K37A)* | **1** | **7.5 ± 0.1** | **52 ± 4** | **100** | **-** |
| | 2 | 6.3 ± 0.3 | 1000 | 28 ± 1 | 4 ± 1 |
| *uvrB(ΔβHG)* | **1** | **7.4 ± 0.1** | **188 ± 46** | **100** | **-** |
| | 2 | 5.9 ± 0.4 | 50 ± 18 | 31 ± 5 | 1.2 ± 0.4 |
| *uvrB(Y96A)* | **1** | **7.7 ± 0.1** | **70 ± 12** | **100** | **-** |
| | 2 | 6.6 ± 0.1 | 1000 | 30 ± 2 | 5.2 ± 0.2 |
| *ΔuvrB* /pUvrB | 1 | 8.1 ± 0.1 | 18.1 ± 0.9 | 100 | - |
| | **2** | **5.9 ± 0.2** | **11.1 ± 0.7** | **26 ± 2** | **0.50 ± 0.08** |

### Supplementary Method

#### ***Strain construction***

Strains of *mfd-YPet* expressing mutants of UvrA and UvrB from the chromosome were constructed using scar-less CRISPR-Cas9 assisted λ Red recombination as previously described[3,4]. Briefly, *mfd-YPet* electrocompetent cells were transformed with pKD46[5] and pCas9[3]. The transformants were plated on LB plate containing ampicillin (50 µg per mL) and chloramphenicol (25 µg per mL) and were incubated at 30 °C overnight. As pCas9 is prone to recombination events[3], the resulting colonies were screened with colony PCR to confirm the presence of the full-length Cas9 gene, using primers targeting upstream and downstream of the Cas9 gene in pCas9.

Next, *mfd-YPet* cells harbouring pKD46 and pCas9 (HH438) were made electro-competent. First, the cells were grown at 30 °C with shaking at 200 rpm in 50 mL of LB containing ampicillin (50 µg per mL) and chloramphenicol (25 µg per mL). In these cells, Cas9 was constitutively expressed. The expression of λ Red recombination proteins was induced with 0.2% L-arabinose (w/v) when cells reached a density around 0.4 ($OD_{600}$). When the optical density reached 0.8, cells were pelleted at 4 °C and washed twice with ice-cold water, with an additional wash using 10% glycerol. Finally, aliquots containing 40 µL of cells in microcentrifuge tubes were snap-freezed in liquid nitrogen and stored at -80 °C.

Cas9 endonuclease activities were targeted to the vicinity of the desired point mutations on the *E. coli* chromosome with the help of the guide RNAs. These guide RNAs were designed such that each contains a 20-nt complementary sequence to that on the *E. coli* chromosome 5' of the PAM sequence (5'-NGG). They were expressed from pCRISPR variants (Extended Data Table 2), which we created following protocols from ref.[3].

The point mutations were introduced by recombining the foreign single-stranded DNA (ssDNA) (Extended Data Table 2). The ssDNAs were 80- to 90-nt oligos flanked by about 40-nt of sequence homologies to the *E. coli* chromosome on both sides of the desired mutations. Base changes were selected to be within five bases from the PAM sequence. This region, termed the seed region, is the most critical to Cas9 binding and disruption in this area ensure cells harbouring the point mutations would not become target for Cas9 cutting.

Aliquots of cells expressing pCas9 and λ Red recombination proteins were transformed with 30 ng of pCRISRP variant plasmid and 500 ng of ssDNA. Positives were selected on LB plates containing 50 µg per mL of kanamycin and 25 µg per mL of chloramphenicol at 37 °C, and were screened by colony PCR, and the promoter and gene sequences were verified. Curing of pCRISPR variant plasmids were performed by propagating cells on LB plates for a week at 42 °C. This is critical for subsequent rounds of genome editing.

### Supplementary References


1 Ho, H. N., Zalami, D., Köhler, J., van Oijen, A.M., Ghodke, H. Identifying multiple kinetic populations of DNA binding proteins in live cells using single-molecule fluorescence imaging. *bioRxiv*, doi:https://doi.org/10.1101/509620 (2019).

2 Truglio, J. J. *et al.* Structural basis for DNA recognition and processing by UvrB. *Nat Struct Mol Biol* **13**, 360-364, doi:10.1038/nsmb1072 (2006).

3 Jiang, W., Bikard, D., Cox, D., Zhang, F. & Marraffini, L. A. RNA-guided editing of bacterial genomes using CRISPR-Cas systems. *Nat Biotechnol* **31**, 233-239, doi:10.1038/nbt.2508 (2013).

4 Pyne, M. E., Moo-Young, M., Chung, D. A. & Chou, C. P. Coupling the CRISPR/Cas9 System with Lambda Red Recombineering Enables Simplified Chromosomal Gene Replacement in Escherichia coli. *Appl Environ Microbiol* **81**, 5103-5114, doi:10.1128/AEM.01248-15 (2015).

5 Datsenko, K. A. & Wanner, B. L. One-step inactivation of chromosomal genes in Escherichia coli K-12 using PCR products. *Proc Natl Acad Sci U S A* **97**, 6640-6645, doi:10.1073/pnas.120163297 (2000).
